# Downregulation of Extraembryonic Tension Controls Body Axis Formation in Avian Embryos

**DOI:** 10.1101/2021.02.24.432525

**Authors:** Daniele Kunz, Anfu Wang, Chon U Chan, Robyn H. Pritchard, Wenyu Wang, Filomena Gallo, Charles R. Bradshaw, Elisa Terenzani, Karin H. Müller, Yan Yan Shery Huang, Fengzhu Xiong

## Abstract

Embryonic tissues undergoing shape change draw mechanical input from extraembryonic substrates. In avian eggs, the early blastoderm disk is under the tension of the vitelline membrane (VM). Here we report that the chicken VM characteristically downregulates tension and stiffness to facilitate stage-specific embryo morphogenesis. Experimental relaxation of the VM early in development impairs blastoderm expansion, while maintaining VM tension in later stages resists the convergence of the posterior body causing stalled elongation, failure of neural tube closure, and axis rupture. Biochemical and structural analysis shows that VM weakening is associated with the reduction of outer-layer glycoprotein fibers, which is caused by an increasing albumen pH due to CO_2_ release from the egg. Our results identify a previously unrecognized potential cause of body axis defects through mis-regulation of extraembryonic tissue tension.

## Introduction

One of the major goals of developmental biology is to understand how large-scale changes in tissue shape (morphogenesis) are regulated at the small-scale molecular level. To drive deformation of soft biological matter in space and time, molecular mechanisms need to generate tissue forces and regulate tissue mechanical properties. While emphasis has been placed on how developing tissues self-organize mechanics internally by controlling cell behaviors, the regulatory mechanisms of the mechanical environment in which the tissues develop are much less understood.

During the early development of many animals, the embryo proper is structurally supported by extraembryonic tissues, which are further attached to the components of the egg. These connections allow transmission of materials, signals and forces between the embryo and the egg environment. As embryonic tissues change shape, they may be assisted or resisted physically by the connected structures (Münster et al., 2019). Measuring the mechanical dynamics of these structures and elucidating their regulatory mechanisms is therefore important for our understanding of developmental defects and the engineering of embryoids and organoids.

Avian embryos offer an excellent model for studying the developmental role and regulation of tissue mechanics because of their large size and accessibility. Inside the egg, the early embryo is supported by a multi-layered protein structure enclosing the yolk, known as the vitelline membrane (VM). The layers of the VM mainly consist of networks of glycoproteins, which provide structural integrity and share homologies to some components of the zona pellucida in mammals (Back et al., 1982; Mann, 2008; Rodler et al., 2012). In chicken embryos, the outer rim of the blastoderm stays attached to the VM during early development, allowing tension transmission between the VM and embryonic tissues (New 1959; Bellairs, Boyde, and Heaysman 1969). The disc-shaped blastoderm expands outwards prior to and during the first day after laying (D0, 0-24hrs) through extensive proliferation and cell movements at the blastoderm edge (epiboly, Figure 1A) (Eyal-Giladi and Kochav, 1976; New, 1959; Sheng, 2014).

**Figure 1.**
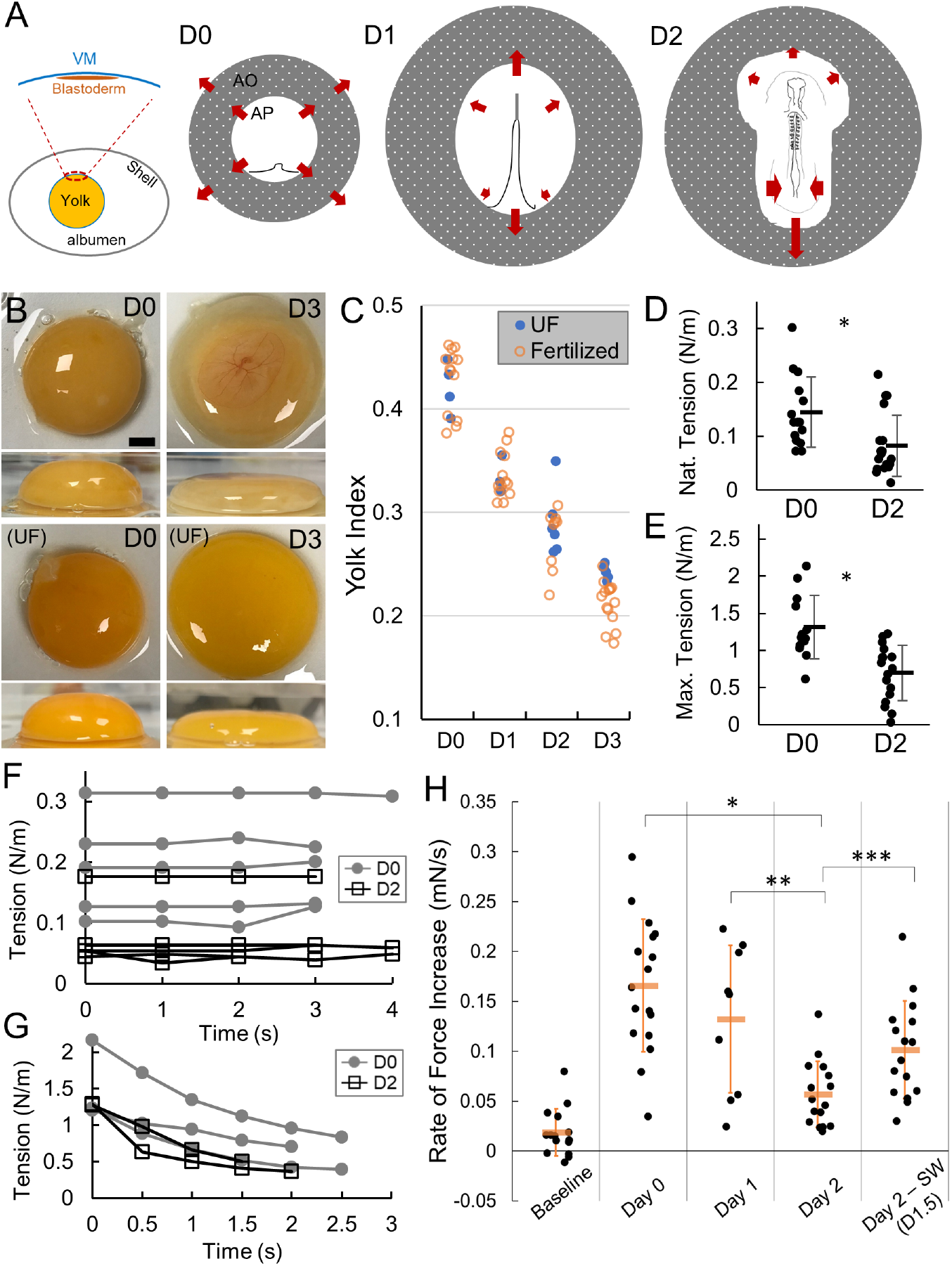
VM tension and strength decrease during early stages of development. A. Diagram of early embryo tissue flow. Left panel shows the blastoderm in relation to the vitelline membrane (VM) and the egg. Right panels illustrate the early shape changes of the blastoderm including area opaca (AO) and area pellucida (AP). Each illustration (anterior to the top) shows the configuration of the blastoderm at approximately the beginning of the labeled incubation day (D0, D1 and D2). D0 involves blastoderm expansion and extension of the primitive streak. D1 and D2 involve continued regression of the streak, tissue convergence and body axis elongation that progress in an anterior to posterior order. Arrows indicate direction of tissue flow. B. Yolk deformation on a flat surface at different stages. Top and side views. UF, unfertilized. Embryo is visible in the D3 image through the red-colored vasculature on the top center of the yolk. Scale bar: 1mm. C. Yolk index (height/diameter) over time. There is no significant difference between UF and fertilized (p>0.05, t-tests). D-E. Native (D) and maximum (E) tensions sustained by the VM. Bars indicate mean+/-SD. *p<0.05, t-tests. F-G. Short-term dynamics of native (F) and maximum tension (G). Each trace represents a single sample. Measurements are taken while the holder is static and the VM is intact. H. VM stiffness measured by the rate of force increase under a stable rate of stretching. Baseline samples do not contain VMs. Day labels indicate the amount of incubation time before the VM samples are measured. Texts following Day labels indicate perturbation type (after the dash) and perturbation start time (in “()”), respectively. For example, “SW (D1.5)” indicates filter sandwich (SW) perturbation started at D1.5. Same for subsequent figures. *,**,*** indicate p<0.05, t-tests.

During the following two days (D1-D2, 24-72hrs), the gastrula embryo converges prominently along the primitive streak and the forming body axis, reversing local tissue flow direction (Figure 1A) (Hamburger and Hamilton, 1951; Rozbicki et al., 2015; Saadaoui et al., 2020; Xiong et al., 2020). Most *ex ovo* culture studies of the chicken embryo found that a stretched VM and its attachment to the blastoderm are necessary for proper long-term development (Bellairs et al., 1967; Chapman et al., 2001; Dugan et al., 1991; New, 1959; Schmitz et al., 2016; Sydow et al., 2017). In the absence of the VM, the blastoderm can only develop when it is supported on a gel substrate (Spratt, 1947) (with posterior axis abnormalities), or when the tension is artificially supplied by inflation through water movement in a “Cornish pasty” culture (Connolly et al., 1995; Nagai et al., 2011). These observations suggest that the extraembryonic physical environment provided by the VM is critical for embryo morphogenesis *in ovo.*

In this study we show that the dynamics of VM mechanics play an essential role in normal development. First, we show that the tension and stiffness of the VM are downregulated during the first 48 hours of incubation. Second, we demonstrate that the early, strong VM is required for blastoderm expansion; while the later, weakened VM is required for the convergent movements of body axis tissues. Third, we show that the changes in tension in the VM result from structural changes of the VM due to the pH increase in the albumen. We show that swapping albumen, lowering pH and reducing CO2 diffusion through the eggshell can biochemically retain a stronger VM, which causes similar body axis phenotypes to those from mechanically increased VM tension. In summary, our results suggest that correct biochemical regulation of the physical properties of the extraembryonic environment is critical for normal embryogenesis.

## Results

### VM weakens during early development

To understand the mechanical dynamics of the VM, we first looked at its effect on the shape of the yolk which it encloses. The yolk index (height/diameter of a separated yolk on a flat surface) is a measure of egg quality in poultry science and reduces characteristically during incubation (Figures 1B-C) (Sauter et al., 1951). The presence of a developing embryo does not alter these dynamics through D3, showing that the cause of change does not come from the embryo itself. At mechanical equilibrium, the tension of the VM balances gravity that drives the spread of the yolk, producing a puddle-like shape.

Because yolks do not show significant changes of volume, mass, or viscosity during these stages (Meuer and Egbers, 1990) (Figure S1A), the VM which wraps the yolk has a stable surface area *in ovo.* When placed on a flat surface, the yolk weight (which is also stable during these stages) stretches the VM. Therefore the lowering yolk index is mainly caused by the VM becoming increasingly stretchable. We modelled the yolk shape by relating the height of the yolk puddle to the VM tension (Figure S1B), which constrains the order of magnitude of tension to 0.1-1 N/m. The VM tension on D2 is predicted to be ~40% of that on D0 from the differences in puddle shape.

To directly measure VM tension, we extracted a piece of VM using windowed filter paper that sticks to the VM (Chapman et al., 2001) and then sandwiched it with a second piece of filter paper to maintain its endogenous stretched state. The sample (kept wet with albumen during the measurements to maintain its *in ovo* chemical environment) is then connected to a weight on one end and vertically suspended on the other. Cutting the horizontal filter paper connections allows the weight to be supported by the VM alone which would reduce the weight reading by a certain amount due to the existing/native tension on the VM (Figures S1C-D). By raising the holder we could increase the tension on the VM until it reaches a breaking point (maximum tension is read before a sharp drop at rupture). We found that, as development proceeds, VM sustains a reduced level of both native and maximum tension (Figures 1D-E), the latter of which is consistent with previous studies using pipette aspiration and texture analyzer experiments on the yolk to evaluate VM tensile strength (Fromm and Matrone, 1962; Kirunda and McKee, 2000). We also noted that the VM behaves elastically at native tension levels (tension is non-zero and remains stable over time, Figure 1F) but viscoelastically when we raised the holder to measure maximum tension (tension drops quickly from high levels while the holder maintains the strain, Figure 1G). While this *ex ovo* method likely introduces some variability during sample preparation, the average magnitudes and differences between D0 and D2 of the VM tension are consistent with our model estimation (Figure S1B). These results show that VM is under tension which decreases from D0 to D2.

To assess the mechanical properties of the VM, we constructed a measurement device that includes a force sensor mounted on a moving stage, a scaffold to hold the VM samples and a side camera (Figure S1E). We used the probe to push perpendicularly on the center of the VM sample attached to filter paper, to stretch the VM while observing its deformation. This method provides a controlled and efficient way to assess VM mechanics as it is compatible with filter paper based *ex ovo* culture of the embryos (Chapman et al., 2001). The force detected by the probe increases as the VM becomes more stretched. We found that D0 and D1 VMs produce higher tension under the same deformation as compared to D2 VMs (Figure 1H). While these deformations likely result in tensions beyond the native levels, the detected differences suggest a sharp reduction of VM stiffness between D0 and D2. Our mechanical model and measurements together demonstrate that VMs weaken characteristically during the early stages of embryonic development.

### VM mechanics affect development of the early embryo

To test the role of VM mechanical changes on embryo morphogenesis, we first subtracted yolk *in ovo* at D0 to cause a reduction of pressure on the VM. The VMs showed visible wrinkles after yolk extraction indicating relaxation of tension. The eggs were then resealed and incubated to D2. This perturbation led to a smaller blastoderm (Figure 2A). Specifically, the area pellucida showed reduced expansion and consequently the body axis length (head to tail) was shorter by >60% (Figures 2A-C). These phenotypes resemble explanted cases where the VM is manually ruffled (Bellairs, Bromham, and Wylie 1967). Control experiments were conducted in which the yolk was extracted and reinjected immediately. These embryos do not show strong defects. These results confirm observations in previous studies (Bellairs et al., 1969; New, 1959) that the early VM needs to be under tension for proper blastoderm expansion.

**Figure 2.**
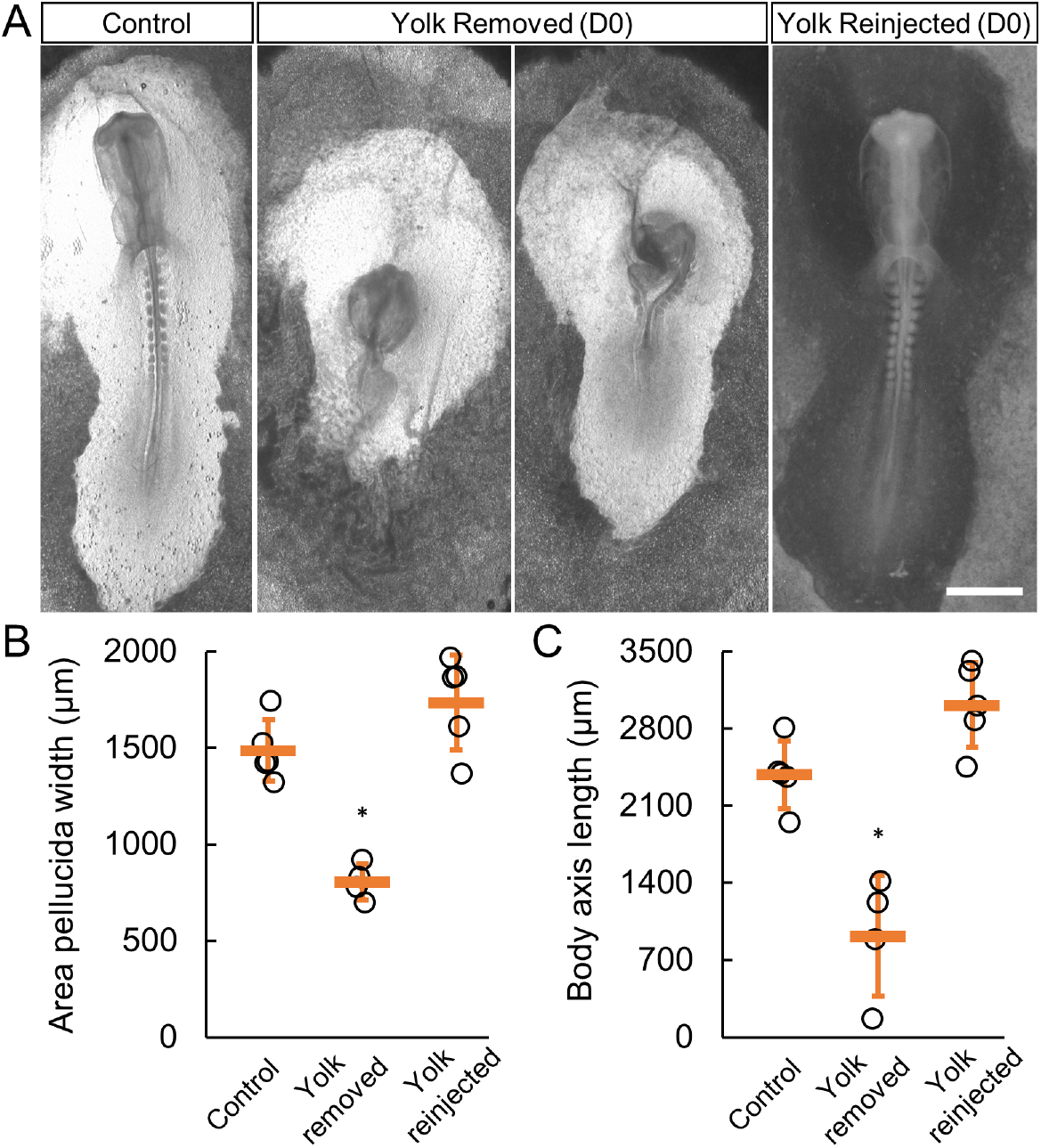
D0 higher VM tension is required for blastoderm expansion. A. Representative embryo phenotypes (D1.5) following yolk extraction and reinjection. Scale bar: 1mm. 2.5ml yolk (~15% V/V) was removed by a syringe with a thin needle through an open window on the eggshell. Wrinkles of VM could be observed after yolk extraction. Eggs with no yolk-leaking were then resealed with tape and incubated. B-C. Width of the area pellucida measured on the posterior end of the body axis (B) and body axis length (head to tail, C) after yolk perturbations. * indicates p<0.01, t-tests.

Conversely, we tested the effect of preventing the observed reduction in tension between D1 and D2 by sandwiching the normal *ex ovo* culture (Chapman et al., 2001) on day 1.5 with an additional piece of windowed filter paper (36-40hrs, Figure S2A). At this stage, the embryo is undergoing gastrulation and body axis formation. The sandwich operation limits the ability of the VM and extraembryonic tissues to move or stretch, resulting in an effectively stiffer tissue environment that is mechanically more similar to D1 rather than the actual stage of D2 (Figure 1H). Under this condition, tissue tension is expected to rise as the large-scale convergence movements underway during these stages pull on the extraembryonic tissues and the VM. To confirm that the embryonic tissues are under a higher tension in sandwiched cultures, we performed surgical cuts through the endoderm and ectoderm lateral to the body axis (Figures S2B-C). The wounds open wider and more persistently in sandwiched embryos after the initial cut but not in control embryos, indicating a higher level of tension in the tissues.

Under enhanced tension, the sandwiched embryos start to show delay of convergence and elongation of the paraxial mesoderm and the neural tube in the posterior body axis (Figures 3A-E; Movies S1-3). Prolonging this condition to ~6hrs results in widening of the axis structures and cases of tissue rupturing (~30%). The ruptures initiate in the posterior midline progenitor region or within the posterior paraxial mesoderm (Movie S3), and then propagate into the neural midline or the paraxial mesoderm along the anterior-posterior axis. These rupture-initiating areas are known to have high cell motility and a reduced level of cell-cell coupling which may underlie their mechanical weakness under tension (Mongera et al., 2019; Xiong et al., 2020). In contrast, in the anterior body, the stiffer somites and neural tubes show a smaller degree of widening under prolonged sandwich culture and do not initiate rupture.

**Figure 3.**
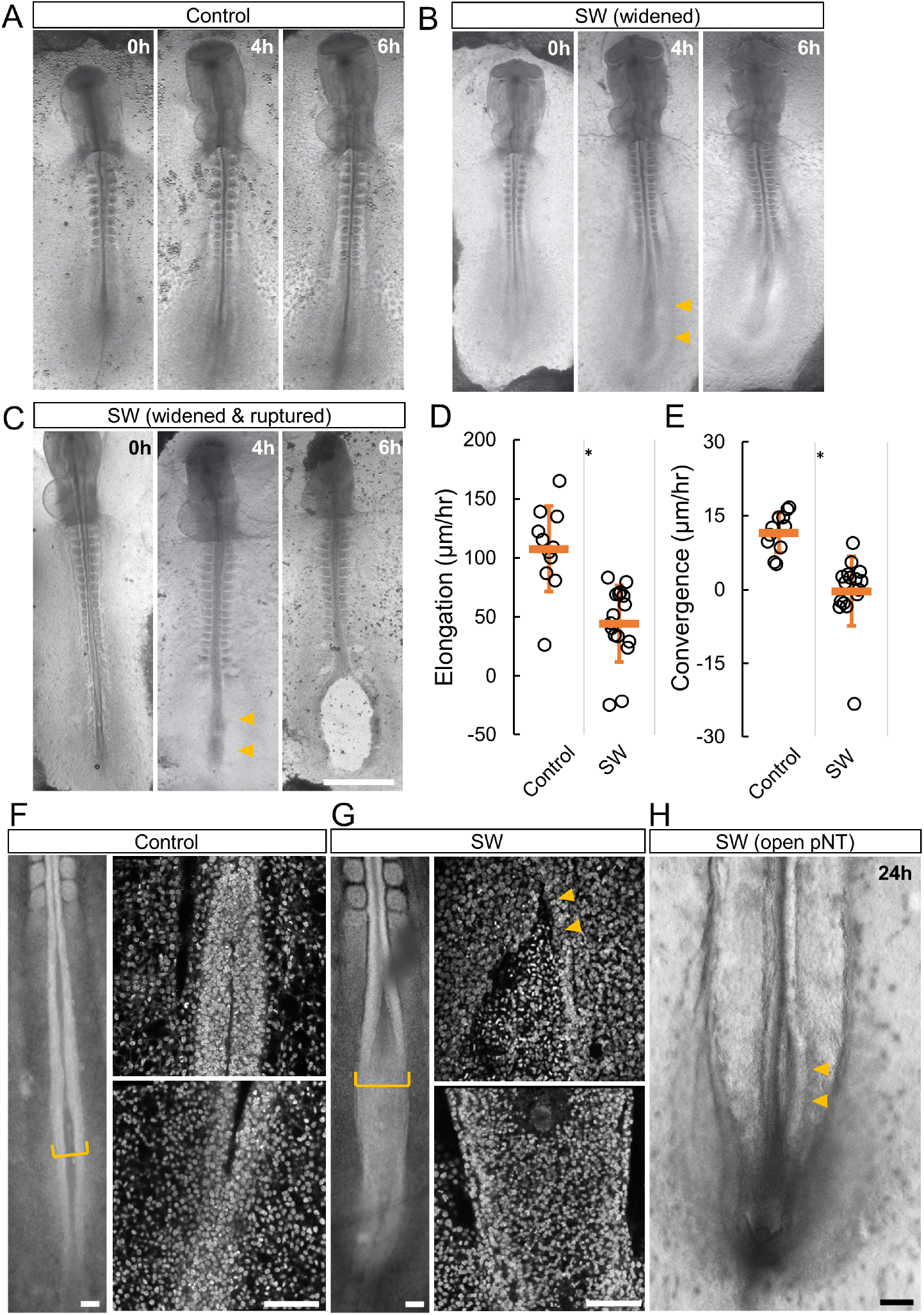
Maintaining higher VM tension in D1-D2 impairs body axis morphogenesis. A-C. Embryo phenotypes (D1.5-D2) following SW. Time stamps indicate hours after the addition of the second filter. Arrowheads indicate the widening posterior body axis. A fraction of SW embryos ruptured under 6 hours (n=11/37). No rupture was observed in controls (n=0/20). Scale bar: 1mm. D-E. Elongation (D) and convergence (E) speeds measured 6 hours after SW in embryos that did not rupture. Some embryos showed a negative convergence speed i.e. widening and/or a negative elongation speed i.e. shortening. Bars indicate mean+/-SD. *p<0.05, t-tests. F-G. Confocal images of the posterior axis comparing Control and SW (nuclear DAPI signal, 6hrs post SW). In the image showing the control embryo the posterior neural tube appeared narrow and mostly closed, while in the SW embryo the posterior neural tube was open and wider. The higher magnification views show the neural tube near the forming somites (top) and the posterior end (bottom), respectively. Scale bars: 100 μm. See also Movies S1-2. H. Open posterior neural tube (pNT) phenotype (n=3/10, arrowheads) in extended SW culture at 24h post SW (D2.5). Scale bar: 100μm.

A majority (~70%) of embryos under the sandwich culture do not rupture but develop a wider and shorter body axis as compared to controls (Movies S1-2; Figures 3B, D-E). This suggests that the mechanical changes in the VM and the tissues caused by this perturbation are not excessive, and is consistent with the mechanical measurements (Figure 1H). Notably, the embryos exhibit a significant delay of neural tube narrowing and, in some cases (~20%), even a reversing of the convergence (resulting in widening). Using confocal microscopy, we found a much flatter, less folded posterior neural plate in sandwiched embryos as compared to the controls (Figures 3F-G). These data show that a higher level of tissue tension impairs the ability of the neural folds to meet and potentially prevents tube closure on the dorsal midline. The increased tension is likely transmitted to the neural epithelium via the non-neural ectoderm, which is known to play a mechanical role in neural tube closure in different systems (Karzbrun et al., 2021; Moon and Xiong, 2021;

Morita et al., 2012; Nikolopoulou et al., 2019; Smith and Schoenwolf, 1997). Indeed, some of the sandwiched embryos show a permanently open posterior neural tube despite developing a normal tail fold (Figure 3H), resulting in a phenotype that resembles mammalian Neural Tube Defects (NTDs) such as spina bifida (Wallingford et al., 2013).

While the sandwich approach as a convenient add-on to the *ex ovo* filter culture method provides a high-throughput way to manipulate tissue tension and allow a detailed analysis of phenotypes, its effect on VM tension is indirect. To increase VM tension directly, we constructed a double-ring culture device that pins down the VM over a liquid chamber connected to an external syringe enabling pressure control (Figures S2D-E) (Sydow et al., 2017). Embryos show strain when the VM is inflated by media injection, consistent with tension transmission between the VM and embryonic tissue (New 1959). Stepwise injection causes the posterior neural folds to widen, and eventually the tissues rupture after multiple rounds of inflation (Figures 4A-B, 10/10). The rupturing points under these conditions also initiate in the posterior midline and paraxial mesoderm. These results mirror the sandwich experiments, suggesting that a moderate increase of VM tension resists convergence while a large increase causes tissue rupture.

**Figure 4.**
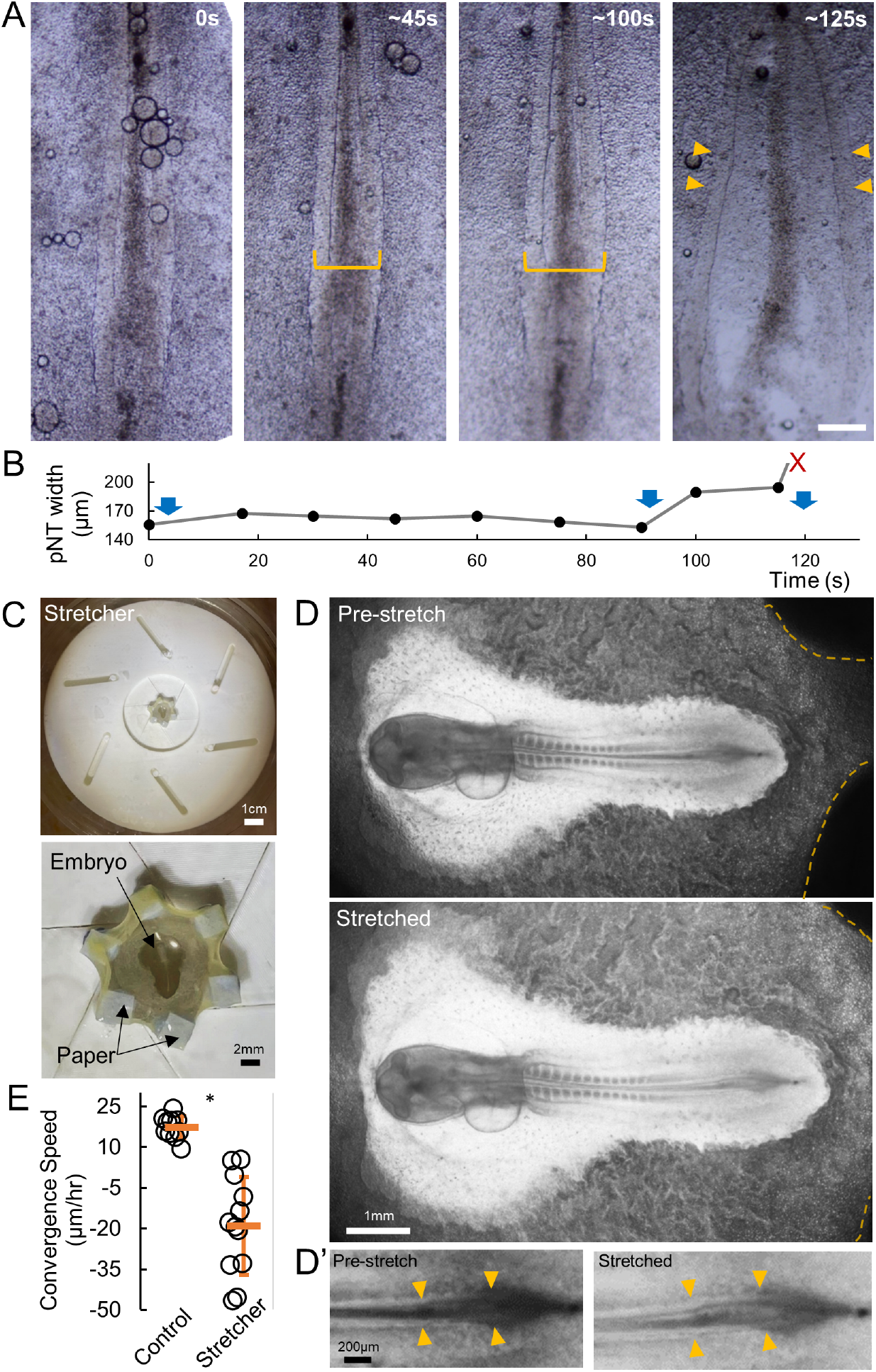
Increasing tension via inflation and stretching causes axis widening. A. Embryo phenotypes following inflation culture. Time stamps indicate seconds after the start of live recording. Yellow brackets measure pNT width as shown in (D). Arrowheads point to the NT tissue after rupture. Posterior body axis widening and/or rupture was observed in all samples tested (n=10/10). Scale bar: 100μm. B. Width of the pNT in inflation culture. Blue arrows indicate times of additional injections of medium to increase inflation and tension. Widening and rupture (red X) follow the injections. C. Rotational stretcher device and example of a mounted embryo. D. Short-term stretching using the stretcher on the VM causes global tissue expansion. The dashed lines mark the contact areas between two of the movable studs and the VM. Under stretching, the embryonic tissues, AP and AO can be seen to enlarge. D’ shows a magnified view of the pNT region where the neural folds can be seen to widen. E. Longer-term convergence speeds measured on the pNT for the ~3-hour window after the embryos were mounted on the stretcher. The controls were not mounted. * indicates p<0.01, t-tests.

The double-ring method works on short timescales and introduces increased pressure on the tissues to increase tension. To distinguish between tension versus pressure as direct causes of the phenotypes, we designed another alternative approach – equiaxial stretcher – to alter VM tension. Similar to a camera shutter, the stretcher uses rotational movement of the blades to generate center to periphery movement, and when attached to the VM via filter paper discs, modulates VM tension by changing VM strain (Figure 4C). We found that embryos loaded on the stretcher show global tissue expansion (Figure 4D). This confirms that tension on the VM is felt throughout the embryo, as only the VM is in contact with the stretcher device, unlike in the sandwich experiments where extraembryonic tissues are also directly affected in addition to the VM. In particular, the posterior body axis tissues (in the example in Figure 4D’, the pNT) shows widening under increased tension. The strain introduced by the stretcher is stable and allows culturing of the embryo (along with the stretcher) over hours, matching the timescale of morphogenesis. We found that the pNTs in embryos mounted on the stretcher show much slower convergence (Figure 4E). These phenotypes are similar to the sandwich and inflation experiments. Further increasing the rotation of the stretcher blades also leads to tissue ruptures. These results show that maintaining VM tension at a higher level is sufficient to disrupt axis convergence in the absence of direct perturbations on the extraembryonic tissues (sandwich method) or pressure (double ring method).

While each approach of tension manipulation has its caveats, they are consistent in producing axis-widening phenotypes that show significant delay in convergence and in some cases ruptures as tension further increases. These results are consistent with the hypothesis that a lowered VM tension is required for normal posterior body morphogenesis between D1 and D2. Our mechanical approaches take advantage of the ease of access to the VM in the *ex ovo* embryo culture. To find methods that prevent tension decrease *in ovo* to further test our hypothesis, a biochemical understanding of VM weakening is required.

### Albumen pH increase due to CO_2_ diffusion causes VM protein loss leading to mechanical weakening

To identify the mechanism of VM mechanical change, we performed electron microscopy to examine the structural changes in the VM during egg incubation. We found a major reduction of protein density in the outer layer of D2 samples as compared to D0 (Figures 5A, S3A). This is consistent with an overall loss of proteins including glycoproteins on the VM (Figure S3B). We stained for glycoproteins directly on the VM and observed significant reduction of fiber density in D2 VMs as compared to D0 (Figures 5B, S3C). Previous studies have shown that the most drastic chemical change in the egg environment during these stages is an increase in the pH of the albumen from ~7.5 upon egg laying to 9-10 in D2 (Fromm 1967), so we tested the effect of pH on isolated VMs. Indeed, treating D0 VMs in pH 9.3 buffer greatly reduces glycoproteins within 12 hours while the fibers are preserved under pH 7.5 (Figures 5C, S3C). The mechanical weakening of VMs in buffer also progresses faster at the incubation temperature (Figure S3D). An increase of pH is known to weaken the interaction strengths between glycoproteins, which reduces the viscosity of the thick albumen and potentially also affects the protein composition of the adjacent VM outer layer (Cotterill and Winter, 1955). We repeated the biochemical assay testing the interaction between ovomucin (a VM glycoprotein) and lysozyme (another major VM component) and confirmed that the solubility of the mixture increases with pH (Figure 5D). These results show that pH controls VM structural integrity which may underlie its mechanical changes during incubation.

**Figure 5.**
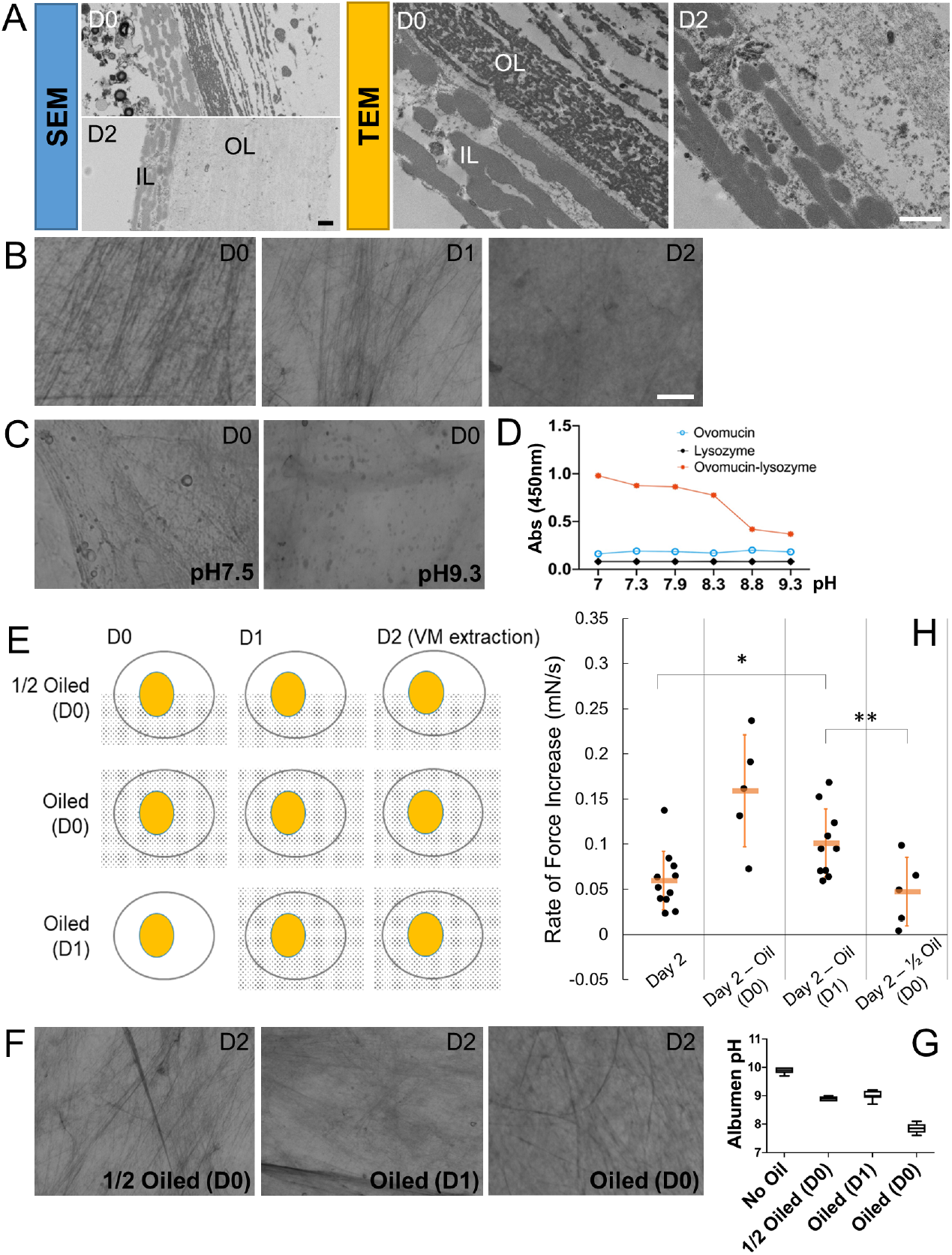
Biochemical changes in the egg environment control VM mechanics. A. Scanning (SEM) and Transmission (TEM) electron microscope cross-sections of the VM. OL, outer layer; IL, inner layer. Scale bar, 1μm. B-C,F. Glycoprotein fibers on the VM stained by the PAS (Periodic acid–Schiff) method. Top right label shows the incubation time when the VM was extracted. Bottom right label shows perturbations. Oil treatments started from the Day indicated in “()”. pH buffer treatments lasted 12 hours. See Figure S3C for additional samples. Scale bar: 10μm. D. Interaction assay between Ovomucin and Lysozyme. Diluted preparations of two components fogged the solution when mixed. The interaction weakened with increasing pH. E. Diagram of oil coating experiments. Dotted patterns indicate oil coating. G. Albumen pH under oil treatments (n=3-5 for each group). Error bars are SD. H. Stiffness measurements of VMs from oiled eggs. *,** indicate p<0.05, t-tests. The method and experimental group labeling in the panel follow those of Figure 1H.

Direct alteration of the albumen pH *in ovo* by adding acids proved challenging, as the albumen becomes murky and embryos do not survive. As an alternative, we performed heterochronic albumen transfer. Eggs incubated to D1 had their albumen removed and replaced with fresh albumen (lower pH) from D0 eggs. These embryos showed a shorter body axis (Figure 6A), consistent with phenotypes of increased VM tension *ex ovo*. The axis lengths in the control transfer (D1 to D1) are indistinguishable from non-operated embryos. For an internal control of body axis length, we compared somite numbers between the experimental groups, which confirmed that embryos were at similar developmental stages (Figure 6A). We also conducted *ex ovo* experiments by culturing the embryos on acidic (pH 6.5) plates and compared these to the normal culture plates (pH 9.0). Embryos on the acidic plates showed slower body axis elongation and convergence speeds (Figures 6B-C). While these results are consistent with tension phenotypes and support a link between pH-controlled VM mechanical properties and embryo development, the disruptive procedures of heterochronic albumen transfer introduce a high degree of variability in phenotypes, and acidic plates may cause posterior axis phenotypes via pathways (Oginuma et al., 2017) other than preventing VM weakening.

**Figure 6.**
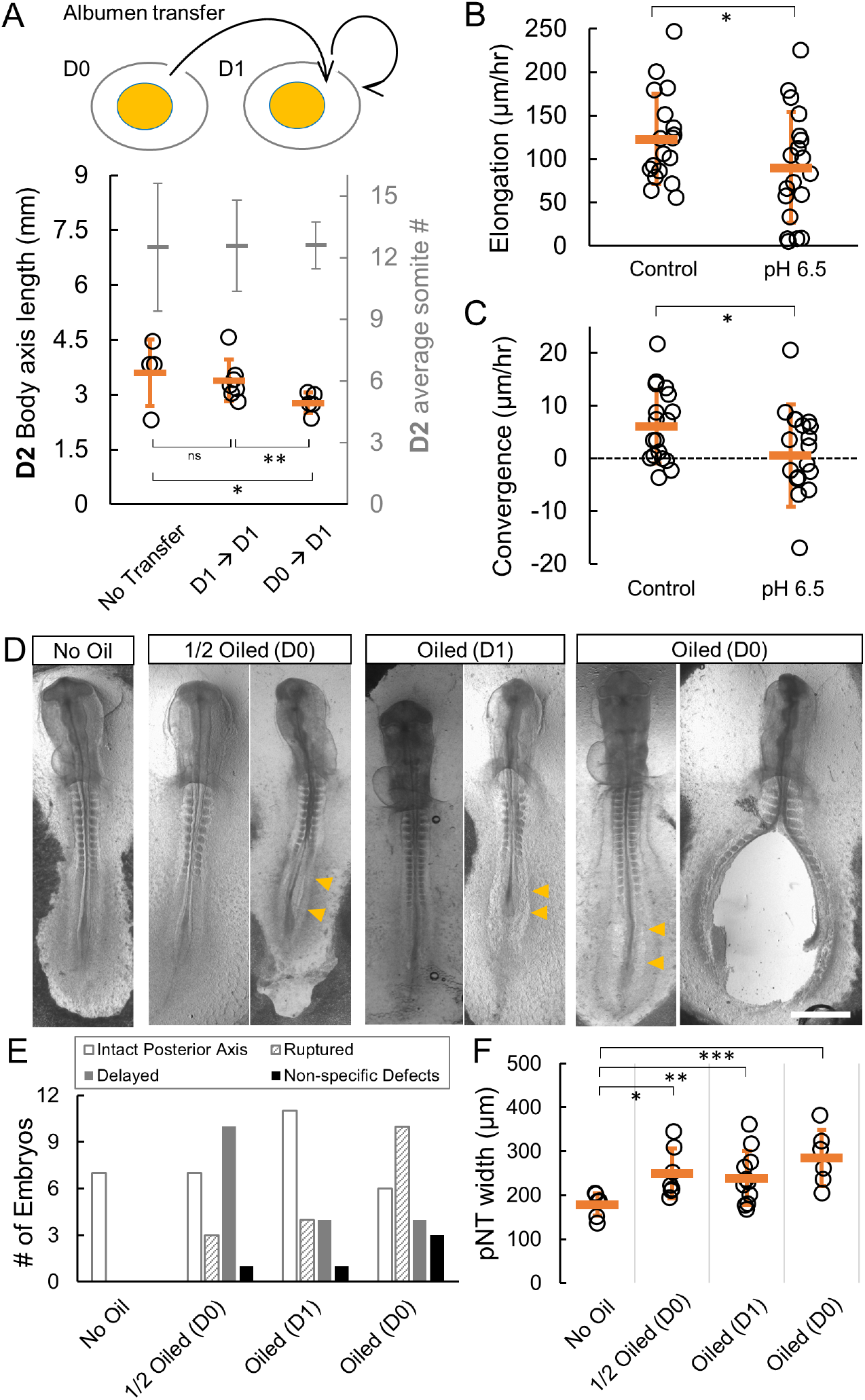
Preventing VM weakening by pH manipulation causes posterior body axis defects. A. Heterochronic albumen transfer including D1→D1 (control) and D0→D1 fresh albumen into D1 eggs (test). Embryos were extracted at D2 and body axis length (from the second somite) was compared. # of somite pairs was counted as an internal control of developmental stage. The test group shows significantly (*,** p<0.05) shorter axis at similar somite stages. B-C. Elongation (B) and Convergence (C) speeds after 6 hours of incubation on acidic (pH 6.5) plates. Some embryos showed a negative convergence speed i.e. widening. Bars indicate mean+/-SD. * indicates p<0.05, t-tests. D. Embryo phenotypes following oil treatments. The embryos were extracted and imaged on D2. Images shown represent the variability of stages and phenotypes in oil treated embryos, summarized in (E). Arrowheads indicate widened posterior body axis tissues. Scale bar: 1mm. E. Summary of phenotypes following oil treatments. Aside from the ruptured embryos, those with 7 or more pairs of somites are categorized as intact posterior axis, embryos with fewer than 7 pairs of somites are categorized as delayed. Embryos with no discernable axis are categorized as non-specific defects. F. Width of the pNT in embryos with intact posterior axis. Bars indicate mean+/-SD. *,**,*** indicate p<0.05, t-tests.

To further reduce the invasiveness of pH manipulation, we used mineral oil to coat the eggshell (Figure 5E), which blocks the release of CO_2_ from air pores that is responsible for the pH increase (Fromm, 1967). This perturbation indeed stalled both the albumen pH increase and glycoprotein fiber reduction (Figures 5F-G). In particular, eggs fully coated in oil at D0 maintain a pH around 7.5 when measured at D2, similar to freshly laid eggs at D0 despite 48 hrs of incubation. Eggs that were half oiled at D0 and eggs fully coated at D1 both showed an intermediate pH at D2, higher than D0 but lower than control eggs.

Importantly, we found that the VMs from fully oiled D0 and D1 eggs retain their respective stiffness (Figure 5H), indicating that VM weakening is stalled by oil treatment. Strikingly, the embryos in the oiled eggs fully recapitulate the spectrum of posterior body axis widening and rupture phenotypes of those observed in the *ex ovo* sandwich, inflation, stretching, pH plate and the *in ovo* albumen transfer experiments (Figures 6D-F). Consistent with maintaining the stiffest VM and highest tension, the eggs fully oiled from D0 produced the most ruptured embryos (Figure 6E). On the other hand, the embryos that developed an intact body axis in these eggs do not show other major defects except the body axis widening phenotypes, suggesting that our oil treatment is not causing systemic failure of development due to the blocking of gas exchange. This is consistent with the low gas exchange requirement of embryos in these early stages (Sadler et al., 1954). Together, these data support the hypothesis that an increase in albumen pH mediated by CO_2_ escape causes VM structural decay during incubation, leading to the mechanical weakening of the VM. This process is necessary for the normal development of the posterior embryonic body, linking the biochemical environment of the egg to the morphogenesis of the embryo.

## Discussion

Tissue morphogenesis does not occur in isolation, but in connection to other tissues and the extraembryonic environment. To some extent, tissues can regulate force production against changes in external resistance and promote robust deformation. However, such adaptation has its limits (Zhou et al., 2015). Our results show that, during the early stages of chicken development, tissue tension is regulated by biochemical modifications of the VM, assisting the tissues to achieve correct size and shape. Interestingly, we found very similar embryonic phenotypes under both physical manipulation of tension and biochemical manipulation of the egg environment. The dynamics of changing tension are likely conserved across the avian phylogeny, as similar yolk shape changes are seen in other avian species (Imai, Mowlah, and Saito 1986; Damaziak et al. 2018). The strong early VM allows extraembryonic cells to generate sufficient tension at the blastoderm edge to drive tissue expansion (Bellairs, Boyde, and Heaysman 1969), and the weak late VM allows body axis tissues to converge to the midline against low resistance. The changes of CO_2_ and albumen pH are triggered by incubation, thereby ensuring that the timing of VM weakening aligns with the developmental progress of the embryo. This provides a novel example of how tissue forces can be biochemically regulated for correct development.

In addition to the biochemical modification of the VM, water influx from the albumen to the yolk may contribute to tension regulation (Moran, 1936). The capacity of tissue water transport is seen in chicken embryos in the Cornish pasty culture condition without the VM (Nagai et al., 2011) and may play a conserved role in size (and probably tension) control in mammalian early embryos (Chan et al., 2019). However, in the D0-D2 early-stage chicken embryos studied here, yolk size increase due to water transport is minimal (Meuer and Egbers, 1990) and does not appear to compensate for the tension reduction due to VM weakening. What the tension dynamics are like in different *ex ovo* culture methods remains to be understood. In the classical New’s method (New, 1955), the embryos are mounted ventral-side up on a watch glass containing a pool of albumen and glass rings hold the VM under tension. This approach essentially maintains the VM-albumen local environment and likely allows the VM to relax tension during incubation in a similar manner to when in the egg. The filter paper dish cultures (Chapman et al., 2001) used in this study and by others have a regular pH around 9, mimicking the albumen of D1-2 eggs that have lost the high body level CO_2_, and are therefore an appropriate environment for VM weakening. It is worth noting that submerged sandwich cultures work very well with no tension phenotypes (Schmitz et al., 2016; Voronov and Taber, 2002), this might be explained by our observation that VM weakening happens quickly when it is soaked in water-based buffers, therefore the sandwiches can no longer cause increased tension. Furthermore, sandwich experiments in late D2 embryos in this study do not show major body axis defects, suggesting that VM tension may no longer be important after the posterior body axis convergence stages.

How global mechanical changes in the VM interact with local stress anisotropy generated by patterned tissue behaviors (Saadaoui et al., 2020) remains to be characterized. The convergence of the body axis tissues is of particular interest given that it is an inward tissue flow and is the most strongly affected process under high tissue tension in our experiments. Convergent movement and folding of the neural plate may involve the non-neural ectoderm generating a pushing force at the tissue transition that aids neural fold formation (Moury and Schoenwolf, 1995). One possible explanation for the failure of neural tube closure in our experiments is that increased tension on the VM limits the movement of these ectodermal cells, preventing the bending of the epithelial sheet that is important for proper folding. Alternatively, the non-neural ectoderm maintained under a higher level of tension could resist the contractile forces generated by the neural plate cells through apical constriction and interkinetic nuclear migration (Nikolopoulou et al., 2017). This would reduce tissue deformation while increasing the load on the cell junctions across the epithelial sheet, making it more likely for tissue ruptures to occur. In either scenario, the requirement of proper tension for tissue convergence raises possible mechanisms with which the neural plate and the non-neural ectoderm actively regulate tissue tension in response to environmental changes. Such mechanisms may include junctional remodeling and cell divisions, which can be tested in future studies using our platform.

The widened/open posterior neural tube in the more mildly perturbed embryos in our study mirrors certain neural tube defects that occur in ~1/1000 human newborns (Fineman et al., 1986; Wallingford et al., 2013). Our results therefore suggest a novel potential cause of neural tube defects through extraembryonic tension mis-regulation. Human embryos at the comparable stages have a similar disc geometry (O’Rahilly and Müller, 2007) and may be under tension through connections with the trophectoderm. The tension dynamics and regulatory mechanisms for the human blastoderm remain unknown and can potentially be studied using *in vitro* cultures or embryoids derived from stem cells (Deglincerti et al., 2016; Karzbrun et al., 2021; Moris et al., 2020). Our results and tools in the avian model enable further studies of genetic and cellular changes under tissue tension, which will shed light on the causes of body axis defects in this critical stage of human development.

## Acknowledgement

The authors thank J. Rees, D. Shah, G. Wu, A. Sossick, P. Williamson, A. Downie, N. Smith, C. Jiggins, G. Sheng and members of the Xiong lab for reagents, technical assistance, and comments. This study is supported by a Wellcome Trust / Royal Society Sir Henry Dale Fellowship (215439/Z/19/Z) to F.X.

## Author Contributions

D.K., A.W. and F.X. designed and performed the experiments, analyzed the data and wrote the manuscript. C.U.C, C.R.B. and E.T. contributed to the ring culture device; R.H.P., *W.W.* and Y.Y.S.H. contributed to the mechanical probe system. F.G. and K.H.M contributed to electron microscopy, which was performed using the facilities of the Cambridge Advanced Imaging Centre (CAIC). The authors declare no competing interest.

## Materials and Methods

### Avian eggs and incubation

Wild type fertilized chicken eggs were supplied by MedEgg Inc. Unfertilized eggs of similar sizes were obtained at Tesco. Eggs were kept in a monitored 15°C fridge for storage and 37.5°C ~45% humidity egg incubators (Brinsea) for incubation. The eggs were incubated within 2 weeks of receipt. No animal protocol was required for the embryonic stages studied under the UK Animals (Scientific Procedures) Act 1986 for incubation of chicken eggs under two weeks. To measure the yolk shape, eggs were opened into a petri dish and both thin and thick albumen was removed carefully with a Pasteur pipette. A top and a side photo of the yolk was taken within 1 minute of albumen clearing, and within 5 minutes of egg opening. The yolk puddle shape reached stability within seconds.

### Imaging

Snapshots of the embryos were taken at different time points with a stereomicroscope (MZ10F, Leica). For timelapse imaging, cultured embryos were transferred to a glass bottom dish (VWR) with a thin layer of culture medium (200μm). Live imaging was performed using a Zeiss Axio Observer 7 microscope equipped with a motorized and heated stage using a 5x objective. Tiles were used to cover the whole embryo and stitched in Zen software to produce movies. For confocal imaging, embryos were first incubated for 6h (control or after filter sandwich placement) and then fixed in 4% paraformaldehyde (PFA) at room temperature for 30 minutes. The nuclei were stained using DAPI (Sigma) 1/2000 for 10min, followed by 2 washes in PBS. Embryos were then mounted on glass bottom dishes. The posterior body axis area of the samples was imaged using a laser scanning confocal microscope with a 40X objective (TCS SP5; Leica).

### Scale measurement of VM tension

Incubated eggs were opened into petri dishes and the top center of the yolk area was cleaned gently to remove the albumen. A strip of filter paper (Whatman) with two identically cut windows of 2cm-wide was folded in the middle to create a loop for a holding pipette mounted on a manual stage. The VM was first attached to one side of the filter paper and cut from the yolk. Then the other side was folded over to fix the VM piece in place. A clip was attached to the open end of the fold to stabilize the mounting as well as serving as the weight on the sample. The whole assembly wais placed inside a digital balance scale (Fisher Scientific). The window has two small triangular cuts in the middle to allow easy release by cutting of the remaining filter paper supporting the weight. This also minimizes VM side deformation caused by the retraction at the cutting site due to tension. After cutting the VM suspends the weight alone. The scale reading of the weight reduces as the holder is raised in a few seconds until the VM breaks, when the reading returns to the maximum. The tension is measured as the force difference over the width of the VM (2cm). See also Figures S1C-D.

### Mechanical probe system

A home-built system was used to measure the mechanical properties of the VM. The system consists of a manual stage (Drummond) with a home-made sample holder, a self-illuminating miniature microscope camera (Kern), and a force sensor (UF1 Isometric, LCMSystems) on a motorized linear translation stage (Thorlabs) connected to a data acquisition system (DAQ, National instruments). See also Figure S1E. The holder uses screws and custom-printed panels to fix the VM samples mounted on filter papers in place, allowing the system to accept samples prepared according to the embryo culture protocol (see the section below). The sensor data and the stage are controlled by a custom program created in LabView (National Instruments). During a measurement run, the VM sample was first mounted on the holder and the holder was then fixed on the manual stage which allows fine positioning of the VM sample to align perpendicularly to the force sensor. The sensor then started logging the measurements. Next the sensor moved towards the VM sample at a constant speed of 0.1mm/s while the approach, contact and VM deformation were recorded from the side by the camera. The pushing phase lasted 40s with contact of VM generally occurring between 5s and 15s. After holding at maximum deformation for 10-15s, the probe was retracted at the same speed.

### Embryo dish culture and filter sandwich

Embryos were staged following the Hamburger and Hamilton (HH) table (Hamburger and Hamilton, 1951). Chicken embryos were extracted for *ex ovo* culture using the Early Chick (EC) culture system (Chapman et al., 2001). For D1.5 controls, the embryos are mostly between stages 9HH and 11HH (approximately 8-somite to 12-somite). To prepare the culture medium, the following formula was used: First, for 100ml of culture media, prepare Part A: 50ml Albumen (beaten for 15min) then supplemented with 0.8ml 20% Glucose and Part B: 0.3g BactoAgar (Sigma) dissolved in 50ml water in a microwave then supplemented with 1.23ml 5M NaCl. Next, equilibrate part A and part B separately in a 55°C water bath. Finally, mix thoroughly and pipette 2ml each to 3.5cm petri dishes before gelation. To make pH 6.5 culture plates, 6M HCl (hydrochloric acid) was used to lower the pH of the A+B mixture to 6.5. The pH was measured using indicator strips. To prepare the filter paper mount for the embryo, 2cm x 2cm pieces of filter paper (Whatman) were punched with two adjacent 0.5cm holes. Eggs were opened into a petri dish. The thick albumen was swept aside gently near the top of the yolk around the embryo. The filter paper was then lowered with the embryo showing through the punched holes and pressed onto the VM. The VM was then cut around the filter paper to release the embryo. The samples were further rinsed in PBS to remove excess yolk. The cleaned embryos were placed ventral side up on pre-warmed culture dishes. The embryo cultures were maintained in a slide box lined with wet paper towels in the egg incubators (37.5°C ~45% humidity) except when snapshot imaging (<2min per embryo) was performed. To perform the filter sandwich perturbation, an additional holed filter paper piece was attached to the ventral side of the embryo on the dish, and then wetted by PBS. To perform surgical cuts in the tissue to assess tension, embryos were first incubated for 4h following the sandwich and control preparation described above. A sharp needle is then used to cut through the endoderm and ectoderm lateral to the body axis. After the surgery the embryos were imaged by an upright Kern microscope using recording mode with 10X magnification, focusing on the cut area.

### Double-ring culture and inflation

The double-ring culture system was designed with modifications from a previous study (Sydow et al., 2017). The base and cover were printed using an in-house 3D printer (Ultimaker S5) using BASF Ultrafuse PLA (Inno3D). The Engineering Profile (0.15mm) was used with 20% infill and no supports. A spherical cover glass (22mm, VWR) was attached to the opening of the base ring by nail polish. The system is wetted with prewarmed culture medium (Part A only, see section above) before use. A 3cm X 3cm square of filter paper with a 2cm diameter hole on the center was used to extract the VM and embryo from the egg. The filter culture embryo was then rinsed in PBS to remove excess yolk. Cleaned VM and embryo were placed on top of the cover glass ventral side up in the medium and clamped between the rings with screws. An externally attached syringe through a tunnel under the base ring allows culture medium to be supplied between the cover glass and the VM, inflating the clamped VM and achieving tension control. Inflation was controlled manually by slowly pressing the syringe to deliver ~1ml of medium each time while observing the VM for rupture or leakage. The experiments were conducted on the stage of a stereomicroscope (10x objective, Kern Optics) and video recorded. The running time of each experiment was usually short term (<10 minutes). With the exception of prewarmed medium, no other temperature control method was implemented.

### Equiaxial tension application

A stretcher device resembling a camera iris was designed and made to apply equiaxial tension to the embryo. The stretcher device consists of three major components: a base, a lid, and six blades. The base was a flat disc (diameter: 12cm) with a circular hole (diameter: 4cm) in the middle. In addition, a hexagonal trench was inscribed on the base disc to allow the blades to slide. The lid had the same dimension as the base, that is, a flat 12cm disc with a 4cm hole in the centre. The lid had six evenly spaced grooves arranged along the tangents to the central hole. The shape of the blade is derived from an equilateral triangle, with two angles being sliced off. Each blade had three additional components: two cylindrical studs facing upward and a cuboid protrusion facing downward. The stud at the tip of the blade was the site of attachment to the VM from the dorsal side of the embryo. The computerized models of these components were designed using Blender (https://www.blender.org/). The .STL file of each component was then generated and 3D-printed using an in-house 3D printer (Ultimaker S5). When the device was assembled, the downward facing protrusion of each blade was placed into the hexagonal trench, whereas the upward facing cylindrical stud near the edge of the blade was inserted into the grooves in the lid. As such, when the lid is rotated, it drove the six blades to slide along the trench which caused the size of the central aperture to change. Since the VM is attached to the studs at the tip of each blade through small pieces of filter paper, as the aperture size enlarged, the VM was stretched and hence equiaxial tension was applied to the embryo. In each test, a HH8-12 embryo was cut from the yolk, transferred to the device and allowed to attach to the filter paper pieces glued to the studs. A thin layer of culture gel or albumen was then added to allow longer term culture of the embryo. See Figure 4C for photos of the configuration. The whole device was then transferred into a 15cm petri dish with lid and then placed in the incubator.

### *In ovo* perturbations

<u>Yolk extractions</u>: a small window (~1.5cmX1.5cm) was opened in the eggshell on D0 eggs with a scalpel blade and scissors. Subsequently a fine pipette was used to puncture the VM and extract yolk from the egg (~2.5 ml). For control embryos ~2.5ml of yolk was removed and immediately reinjected. After yolk extraction the eggs were sealed with tape and incubated at 37.5°C for 38h (D1.5). The embryos were extracted and imaged (Leica MZ10F). Samples that did not heal the VM resulting in yolk collapse would fail to develop embryos and are excluded from analysis. <u>Oil treatment</u> (adapted from (Fromm, 1967)): Eggs were dipped in mineral oil (Sigma) and wiped clean of excess oil and returned to the incubators. The oil stayed on the eggshell pores stably over extended time (48h maximum in this study) as observed under bright light. For <u>heterochronic albumen transfer</u>, a small window (~1.5cmx1.5cm) was opened in the eggshell on D1 eggs and the albumen was extracted slowly using a pipette without damaging the yolk. Fresh albumen (D0) was then injected into the eggs (D1). For the control samples the albumen (D1) was removed and reinjected. After the manipulation the eggs were sealed with tape and incubated at 37.5°C for ~24h to reach D2. The embryos were then extracted and imaged (Leica MZ10F).

### VM protein extraction and SDS-PAGE

~8 eggs per incubation time (0h; 24h; 36h; 48h; 54h) were opened into petri dishes and the VMs were cut open, removed from the dish and washed in PBS. The obtained VMs were dried in a SpeedVac. Proteins were extracted from the samples by using a buffer consisting of 50 mM Tris-HCl (pH 8.0), 10% glycerol, 2% sodium dodecyl sulfate (SDS), 25 mM ethylenediaminetetraacetic acid, and protease inhibitor cocktail (Sigma-Aldrich). Samples were incubated overnight under constant stirring at room temperature. After this, the samples were centrifuged (12000g, 30 min, 4°C), the supernatant was collected, and the concentration of proteins was determined using the Lowry method. Sodium dodecyl sulfate polyacrylamide gel electrophoresis (SDS-PAGE) was carried out using 1.0 mm thickness of both running gel (12%) and stacking gel (5%) in the presence of SDS. The electrophoresis was run at 8 V/cm in a Tris-HCl buffer system using a Vertical Biorad Mini attached to a Power Pac Basic electrophoresis apparatus. The samples were dissolved in 0.2mol/L Tris-HCl buffer (pH 8.0) containing SDS (5g/L) to a final concentration of 100μg of protein/lane. Protein solutions (10μl) along with a molecular weight ladder (260kD, Thermo Scientific) were loaded onto the gel and stained with colloidal Coomassie blue (Colloidal Blue Stain Kit, Invitrogen) or with Pierce^™^ Glycoprotein Staining Kit (Thermo Scientific) after separation.

### Glycoprotein staining of the vitelline membrane

To examine the glycoprotein fibers on the VM, the samples were mounted on a cover slide and stained using Pierce™ Glycoprotein Saining Kit (Thermo Scientific), with PBS buffer wash to prevent drying. The stained VMs were examined microscopically at 100x (Kern Optics). To test the effect of pH changes on the VM glycoproteins, eggs without incubation (D0) were used to extract the VM. The samples were soaked in pH buffers of 7.5 and 9.3. The buffers were made with 0.1M monopotassium phosphate and 0.05M borax (pH 7.9 to 9.5) and 0.05M phosphate buffer (Hawthorne, 1950). No adjustments were made for ionic strength. The VM samples were incubated overnight (12h) at room temperature (~25°C) or in the incubator (37.5°C).

### Lysozyme-ovomucin interaction under different pH

Ovomucin was prepared as described (Omana and Wu, 2009). Albumen was diluted with 5 volumes of 0.1 mol/L NaCl solution and stirred gently for 30min using a magnetic stirrer. Subsequently, the pH of the mixture was adjusted to approximately 6 using 1 mol/L HCl. The dispersion was kept overnight at 4°C and centrifuged at 15,000 g for 20min at 4°C. The precipitate was suspended in 0.5 mol/L NaCl solution and then kept at 4°C for 6h. After another centrifugation at 15,000 g for 20 min at 4°C, the precipitate (containing ovomucin) was separated from the supernatant. The preparation was analyzed with SDS-PAGE to check for ovomucin. A few other protein bands were found to remain in the preparation but not expected to affect the following assay. The interaction between lysozyme and ovomucin was tested by turbidity as described (Cotterill and Winter, 1955). Tubes containing the ovomucin preparation were diluted 1:2 with pH 7.0 to 9.3 buffers before the addition of purified lysozyme (2ug/ml, Sigma). The contents were then mixed and incubated at 37°C for 1 hour. The absorbance was measured at 450nm by a microplate reader (Perkin Elmer 1420).

### Sample preparation for electron microscopy

Eggs from different time incubation (0h; 48h; 54h) were used for VM extraction. Samples were fixed in a fixative (2%glutaraldehyde/2% formaldehyde in 0.05 M sodium cacodylate buffer pH 7.4 containing 2 mM calcium chloride) overnight at 4°C. After washing 5x with 0.05M sodium cacodylate buffer at pH 7.4, samples were osmicated (1% osmium tetroxide, 1.5% potassium ferricyanide, 0.05M sodium cacodylate buffer at pH 7.4) for 3 days at 4°C. After washing 5x in DIW (deionised water), samples were treated with 0.1% (w/v) thiocarbohydrazide/DIW for 20 minutes at room temperature in the dark. After washing 5x in DIW, samples were osmicated a second time for 1 hour at RT (2% osmium tetroxide/DIW). After washing 5x in DIW, samples were block-stained with uranyl acetate (2% uranyl acetate in 0.05 M maleate buffer at pH 5.5) for 3 days at 4°C. Samples were washed 5x in DIW and then dehydrated in a graded series of ethanol (50%/70%/95%/100%/100% dry), 100% dry acetone and 100% dry acetonitrile, 3x in each for at least 5 min. Samples were infiltrated with a 50/50 mixture of 100% dry acetonitrile/Quetol resin (without BDMA) overnight, followed by 3 days in 100% Quetol (without BDMA). Then, the sample was infiltrated for 5 days in 100% Quetol resin with BDMA, exchanging the resin each day. The Quetol resin mixture is: 12 g Quetol 651, 15.7 g NSA, 5.7 g MNA and 0.5 g BDMA (all from TAAB). Samples were placed in embedding moulds and cured at 60°C for 3 days.

### TEM and SEM

For TEM (transmission electron microscopy) imaging, sections (70nm thickness) were prepared in an ultramicrotome (Leica Ultracut) and placed on 300 mesh bare copper grids. Samples were viewed in a Tecnai G2 TEM (FEI/ThermoFisher) run at 200 keV accelerating voltage using a 20μm objective aperture to improve contrast; images were acquired using an AMT digital camera. For SEM (scanning electron microscopy) imaging, thin sections were placed on Melinex (TAAB) plastic coverslips and allowed to air dry. Coverslips were mounted on aluminium SEM stubs using conductive carbon tabs (TAAB) and the edges of the slides were painted with conductive silver paint (TAAB). Then, samples were sputter coated with 30 nm carbon using a Quorum Q150 T E carbon coater. Samples were imaged in a Verios 460 scanning electron microscope (FEI/Thermofisher) at 4 keV accelerating voltage and 0.2nA probe current in backscatter mode using the concentric backscatter detector (CBS) in field-free mode at a working distance of 3.5-4mm; 1536×1024 pixel resolution, 3μs dwell time, 4 line integrations. Stitched maps were acquired using FEI MAPS software using the default stitching profile and 10% image overlap.

### Data analysis

Images and movies were processed by Fiji (NIH) and Powerpoint (Microsoft) for measurements. Scale bars were first set with control images with objects of known sizes. For elongation speed, the distance between a fixed somite pair (usually somite 3 or 4) and the posterior end of the body axis was taken. For posterior neural tube (pNT) width and convergence speed, the distance between the outer boundary of the neural tube walls was taken. For mechanical probe data, the raw trace (TDMS file, LabView) of each measurement containing the position of the probe and the force data over time was loaded into Matlab using custom code including a TDMS converter (Brad Humphreys, https://github.com/humphreysb/ConvertTDMS). The traces were examined alongside their corresponding live movies to exclude situations of early VM rupture and when probes hit the holder structure. The force curve of the admitted traces was first adjusted with their own baseline before probe movement started. The high frequency data points were then averaged to one value per second. The average rate of force increase of a 20s duration over the approximately linear phase of force curve was used to measure the stiffness of the samples.

## Supplementary Information

**Figure S1.**
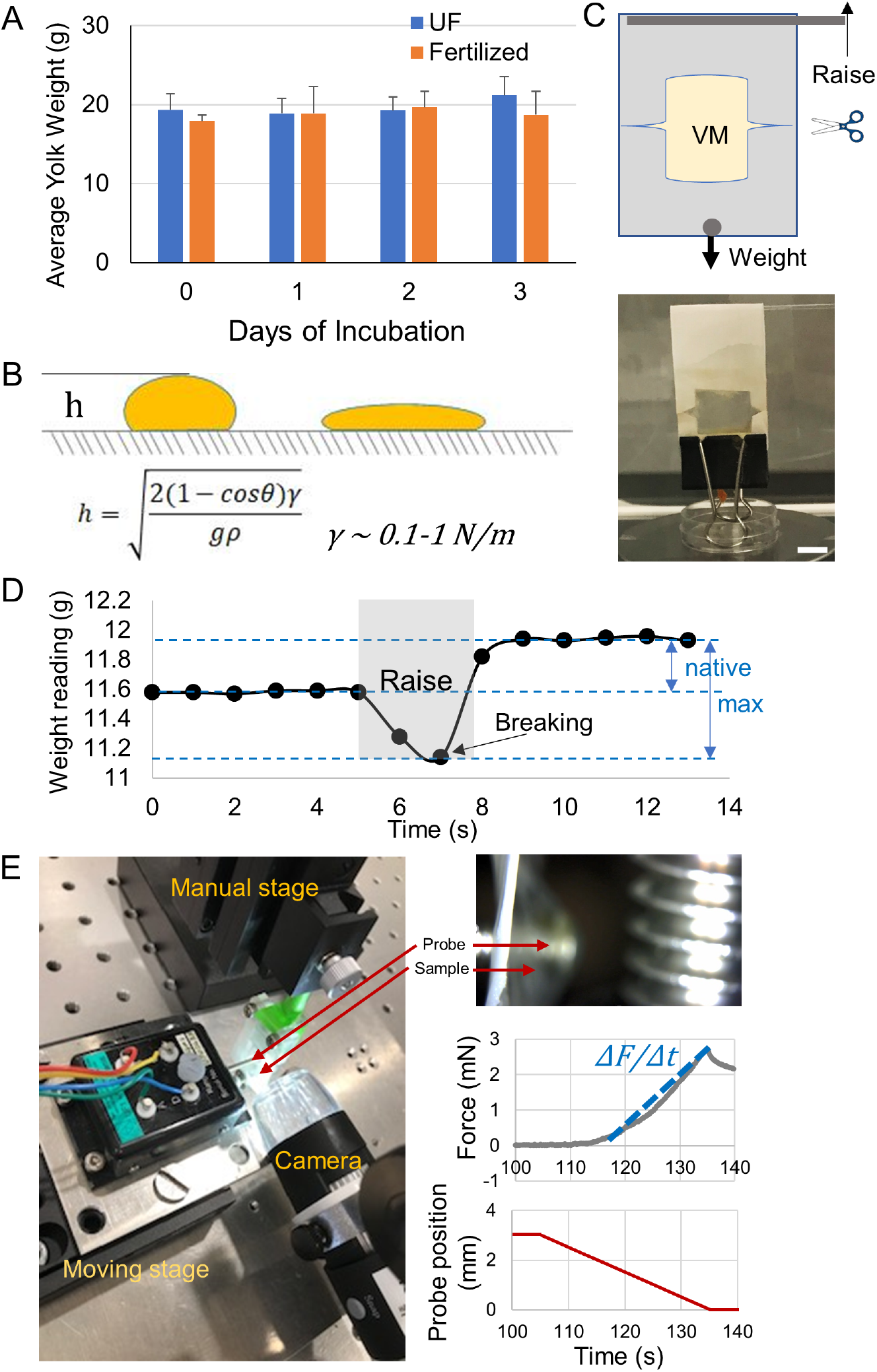
Physical changes of the yolk and the VM. A. Average yolk weight. Albumen was removed as much as possible for each yolk with a Pasteur pipette. n=3-6 for unfertilized eggs (UF); n=8-14 for fertilized eggs. Error bars show +SD. The weights were similar (p>0.05, t-tests). B. Simple droplet model of the yolk shape (de Gennes et al., 2013). *h*(height)~1-2×10^-2^m, *ρ*(density)~1×10^3^kg/m^3^, *g*(gravitational acceleration)~10m/s^2^, *θ*(contact angle)>90°. Using our height data (Figure 1C), The VM tension was estimated to be ~1.68 N/m on D0 and ~0.67 N/m on D2. C. Tension measurement using a scale. Schematic on the left and an example on the right. The VM was sandwiched between two identical sides of a folded piece of windowed filter paper. A clip serving as the weight was attached on the bottom. Small contact points on both sides were designed in the window to minimize side deformation of the VM. After cutting these points the VM suspended the weight and raising the holder increased VM tension. Weight reading was recorded and analyzed in a live video. Scale bar, 1cm. D. Example of a typical raise experiment. The maximum drop point was identified in the video which usually corresponds to the visible breakage starting to appear. This and the pre-raise line were compared to the stable weight to calculate tensions. E. Stiffness measurement using a force probe. The left photo shows the components of the homemade system. Top right shows a measurement in progress from the camera view. The probe is seen to protrude and deform the VM towards the right side away from the filter mount. The plots show an example measurement sequence on a parafilm control. The probe moved at a stable speed and started to detect a force when the sample started to resist deformation. The average rate of increase (mN/s) of a 20s duration over the approximately linear phase of force curve *(ΔF/Δt* in the plot) was used to measure VM resistance.

**Figure S2.**
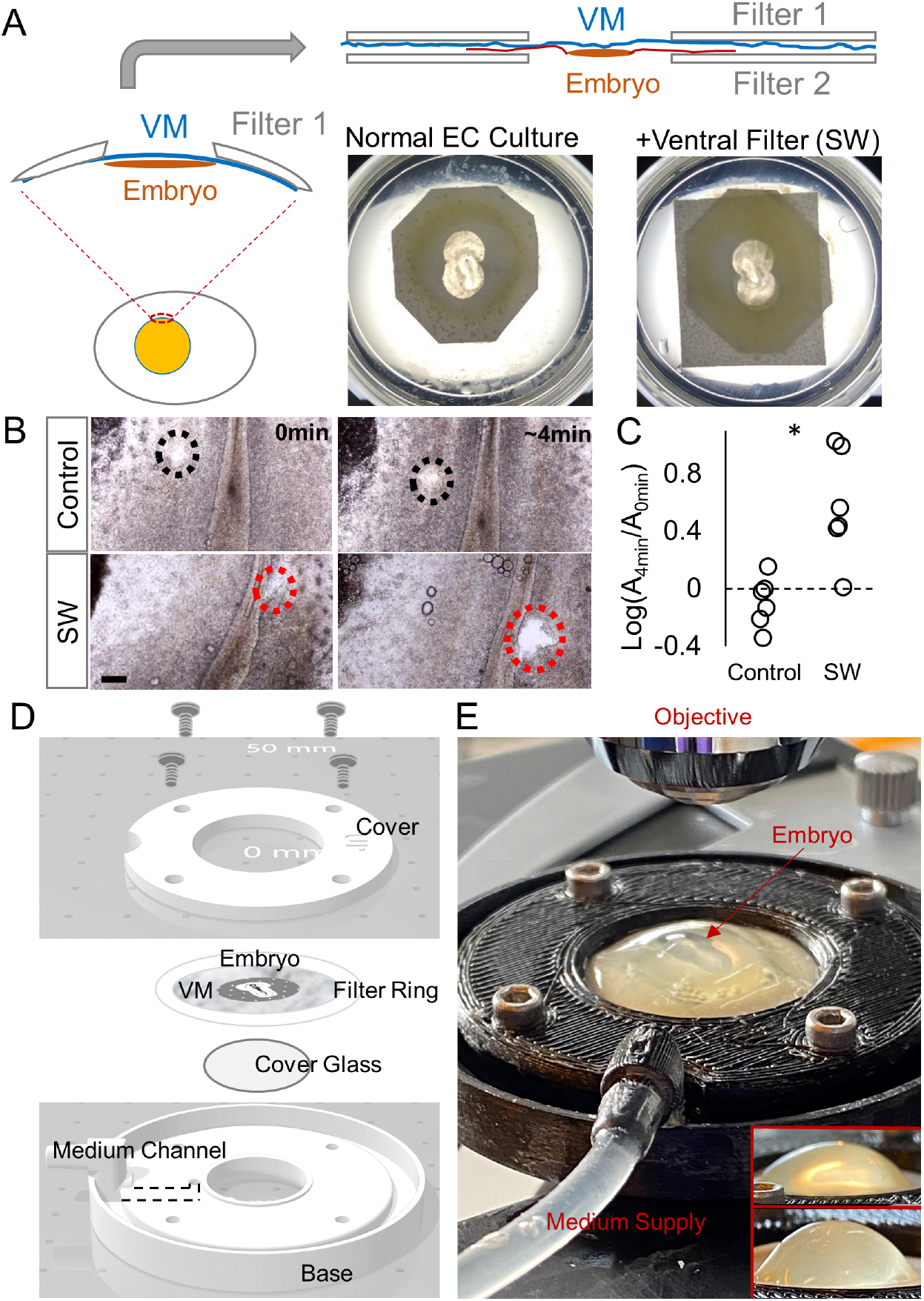
Tension perturbations of the VM. A. Filter sandwich (SW) to maintain tension of the VM. In the normal *ex ovo* culture protocol (Chapman et al., 2001), one piece of windowed filter paper (Filter 1) is attached to the VM and the embryo is cut out and laid down on the culture dish with ventral side up. By placing another filter (Filter 2) on the ventral side, the VM and the distal part of the extraembryonic tissue were sandwiched and fixed in place. Tissues in the window (including the embryo) can continue their movements. B. Wound opening after a needle cut on the embryos. Circles mark the wound site. Scale bar, 100μm. C. Wound size change. Areas at 0min and approximately 4min after cut are compared. Controls show slow opening or closing while SWs show consistent expansion. *p<0.05 (t-test). D-E. VM tension modulation with a double-ring inflation culture device. This system is a new design inspired by (Sydow et al., 2017). The schematic (D) shows the components of the set-up. The photo (E) shows an experiment (Inserts show side views of different degrees of inflation, top is mild, bottom is strong). The cover and base (i.e. the double ring) were 3D printed. Embryo extracted (D1.5) on a filter ring is laid ventral side down on a cover glass which is glued to the opening of the base ring. Screws secure the seal of the sample allowing syringe controlled medium supply to inflate the sample. Samples with damage on the VM causing medium leaking were discarded.

**Figure S3.**
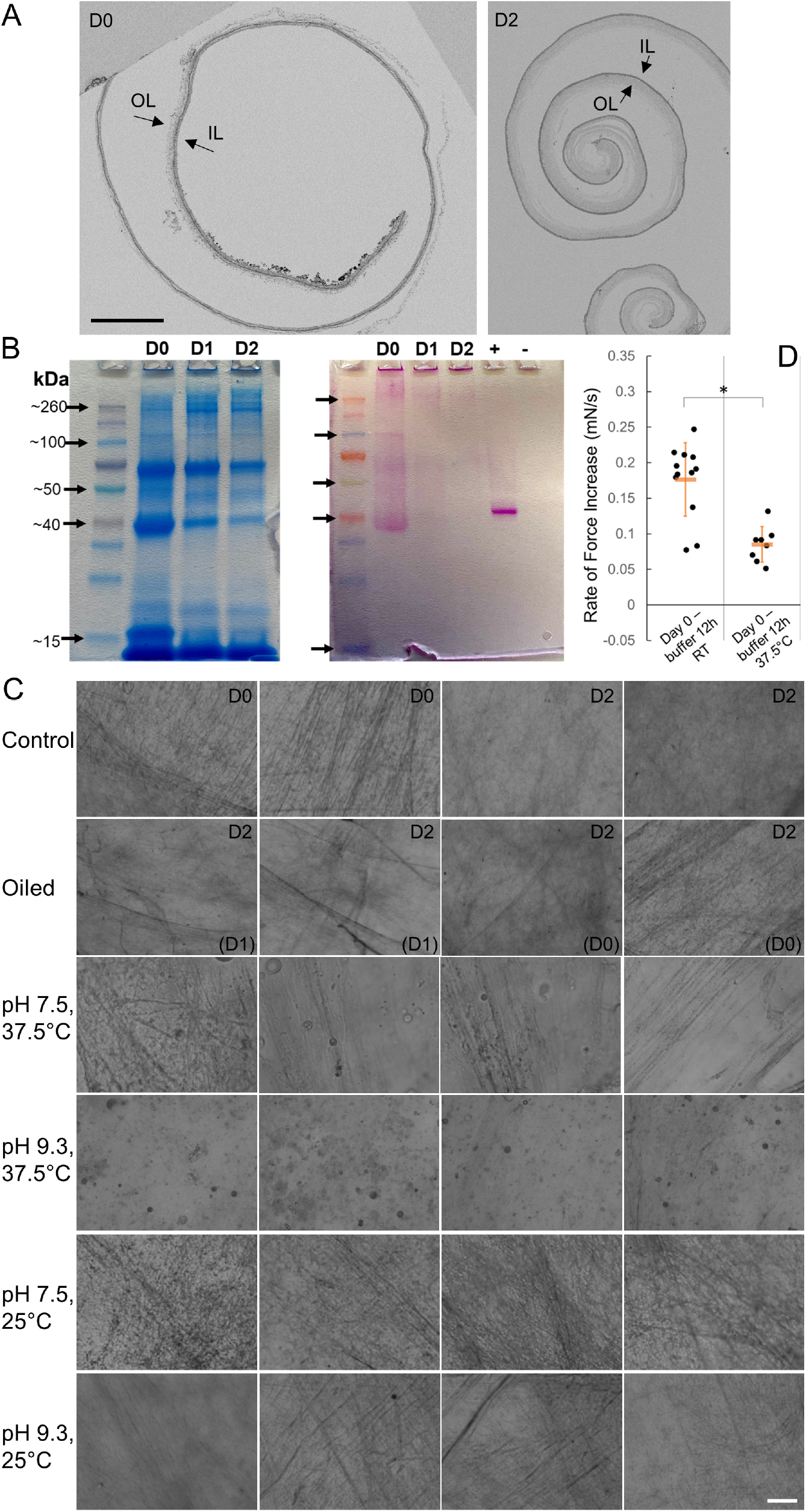
Structural changes of the VM. A. Lower magnification SEM views of the VM cross-sections. OL, outer layer; IL, inner layer. The VM patterns shown in Figure 5A are uniformly found on the samples. Scale bar, 100μm. B. SDS gels of VM protein extraction stained by Coomassie blue (left) and PAS for glycoproteins (right), respectively. C. Glycoprotein fibers on the VM (these are additional examples related to Figures 5B-C,F). Top right label shows the Day when the VM is extracted and fixed for PAS staining. Bottom right label shows the Day when oil treatment starts. For the pH buffer treatment (12 hours) groups, all VMs are D0. Scale bar, 10μm. D. Stiffness measurement of buffer treated VM samples under different temperatures.

**Movies S1-S3. Example timelapse of body axis morphogenesis**

Anterior to the left. S1 is a control embryo, consistent posterior body axis narrowing and elongation are visible. S2 is a SW embryo visibly shorter and wider. S3 is a SW embryo that ruptured. The starting times of the movies are around ~0.5hr after the placing the SW filter taking into account the time need for mounting and timelapse set-up. Scale bars, 500μm.

## Notes

### Competing Interest Statement

The authors have declared no competing interest.

### Summary of Updates

New controls added to Yolk extraction experiments New method/data added to tension control - stretcher New method/data added to biochemical tests - heterochronic albumen transfer Additional panels to supplementary figures for clarity Overall writing and organization improved. Manuscript extended (6 main figures)

## References

Back JF, Bain JM, Vadehra D v., Burley RW. 1982. Proteins of the outer layer of the vitelline membrane of hen’s eggs. Biochimica et Biophysica Acta (BBA)/Protein Structure and Molecular 705:12–19. doi:10.1016/0167-4838(82)90329-6

Bellairs R, Boyde A, Heaysman JEM. 1969. The relationship between the edge of the chick blastoderm and the vitelline membrane. Wilhelm Roux’ Archiv für Entwicklungsmechanik der Organismen 163:113–121. doi:10.1007/BF00579315

Bellairs R, Bromham DR, Wylie CC. 1967. The influence of the area opaca on the development of the young chick embryo. Journal of Embryology and Experimental Morphology 17:195–212.

Chan CJ, Costanzo M, Ruiz-Herrero T, Mönke G, Petrie RJ, Bergert M, Diz-Muñoz A, Mahadevan L, Hiiragi T. 2019. Hydraulic control of mammalian embryo size and cell fate. Nature 571:112–116. doi:10.1038/s41586-019-1309-x

Chapman SC, Collignon JJ, Schoenwolf GC, Lumsden A. 2001. Improved method for chick whole-embryo culture using a filter paper carrier. Developmental Dynamics 220:284–289. doi:10.1002/1097-0177(20010301)220:3<284::AID-DVDY1102>3.0.CO;2-5

Connolly D, McNaugbton LA, Krumlauf R, Cooke J. 1995. Improved in vitro development of the chick embryo using roller-tube culture. Trends in Genetics 11:259–260. doi:10.1016/S0168-9525(00)89070-8

Cotterill OJ, Winter AR. 1955. Egg White Lysozyme. Poultry Science 34:679–686. doi:10.3382/ps.0340679

Damaziak K, Marzec A, Kieliszek M, Buclaw M, Michalczuk M, Niemiec J. 2018. Comparative analysis of structure and strength of vitelline membrane and physical parameters of yolk of ostrich, emu, and greater rhea eggs. Poultry Science 97:1032–1040. doi:10.3382/ps/pex356

de Gennes PG, Brochard-Wyart F, Quéré D. 2013. Capillarity and wetting phenomena: drops, bubbles, pearls, waves. Springer Science & Business Media.

Deglincerti A, Croft GF, Pietila LN, Zernicka-Goetz M, Siggia ED, Brivanlou AH. 2016. Self-organization of the in vitro attached human embryo. Nature 533:251–254. doi:10.1038/nature17948

Dugan JD, Lawton MT, Glaser B, Brem H. 1991. A new technique for explantation and in vitro cultivation of chicken embryos. The Anatomical Record 229:125–128. doi:10.1002/ar.1092290114

Eyal-Giladi H, Kochav S. 1976. From cleavage to primitive streak formation: A complementary normal table and a new look at the first stages of the development of the chick. I. General morphology. Developmental Biology 49:321–337. doi:10.1016/0012-1606(76)90178-0

Fineman RM, Schoenwolf GC, Huff M, Davis PL. 1986. Animal model: Causes of windowing-induced dysmorphogenesis (neural tube defects and early amnion deficit spectrum) in chicken embryos. American Journal of Medical Genetics 25:489–505. doi:10.1002/ajmg.1320250311

Fromm D. 1967. Some Physical and Chemical Changes in the Vitelline Membrane of the Hen’s Egg During Storage. Journal of Food Science 32:52–56. doi:10.1111/j.1365-2621.1967.tb01956.x

Fromm D, Matrone G. 1962. A Rapid Method for Evaluating the Strength of the Vitelline Membrane of the Hen’s Egg Yolk. Poultry Science 41:1516–1521. doi:10.3382/ps.0411516

Hamburger V, Hamilton HL. 1951. A series of normal stages in the development of the chick embryo. Journal of Morphology 88:49–92. doi:10.1002/jmor.1050880104

Hawthorne JR. 1950. The action of egg white lysozyme on ovomucoid and ovomucin. BBA - Biochimica et Biophysica Acta 6:28–35. doi:10.1016/0006-3002(50)90074-6

Imai C, Mowlah A, Saito J. 1986. Storage Stability of Japanese Quail (Coturnix coturnix japonica) Eggs at Room Temperature. Poultry Science 65:474–480. doi:10.3382/ps.0650474

Karzbrun E, Khankhel AH, Megale HC, Glasauer SMK, Wyle Y, Britton G, Warmflash A, Kosik KS, Siggia ED, Shraiman BI, Streichan SJ. 2021. Human neural tube morphogenesis in vitro by geometric constraints. Nature 2021 599:7884 599:268–272. doi:10.1038/s41586-021-04026-9

Kirunda DFK, McKee SR. 2000. Relating quality characteristics of aged eggs and fresh eggs to vitelline membrane strength as determined by a texture analyzer. Poultry Science 79:1189–1193. doi:10.1093/ps/79.8.1189

Mann K. 2008. Proteomic analysis of the chicken egg vitelline membrane. Proteomics 8:2322–2332. doi:10.1002/pmic.200800032

Meuer H-J, Egbers C. 1990. Changes in density and viscosity of chicken egg albumen and yolk during incubation. Journal of Experimental Zoology 255:16–21. doi:10.1002/jez.1402550104

Mongera A, Michaut A, Guillot C, Xiong F, Pourquié O. 2019. Mechanics of anteroposterior axis formation in vertebrates. Annual Review of Cell and Developmental Biology 35:259–283. doi:10.1146/annurev-cellbio-100818-125436

Moon LD, Xiong F. 2021. Mechanics of neural tube morphogenesis. Seminars in Cell & Developmental Biology. doi:10.1016/J.SEMCDB.2021.09.009

Moran T. 1936. Physics of the Hen’s Egg: II. The Bursting Strength of the Vitelline Membrane. Journal of Experimental Biology 13:41–47.

Moris N, Anlas K, van den Brink SC, Alemany A, Schröder J, Ghimire S, Balayo T, van Oudenaarden A, Martinez Arias A. 2020. An in vitro model of early anteroposterior organization during human development. Nature 582:410–415. doi:10.1038/s41586-020-2383-9

Morita H, Kajiura-Kobayashi H, Takagi C, Yamamoto TS, Nonaka S, Ueno N. 2012. Cell Movements of the deep layer of non-neural ectoderm underlie complete neural tube closure in Xenopus. Development 139:1417–1426. doi:10.1242/dev.073239

Moury JD, Schoenwolf GC. 1995. Cooperative model of epithelial shaping and bending during avian neurulation: Autonomous movements of the neural plate, autonomous movements of the epidermis, and interactions in the neural plate/epidermis transition zone. Developmental Dynamics 204:323–337. doi:10.1002/aja.1002040310

Münster S, Jain A, Mietke A, Pavlopoulos A, Grill SW, Tomancak P. 2019. Attachment of the blastoderm to the vitelline envelope affects gastrulation of insects. Nature 568:395–399. doi:10.1038/s41586-019-1044-3

Nagai H, Lin MC, Sheng G. 2011. A modified cornish pasty method for ex ovo culture of the chick embryo. Genesis 49:46–52. doi:10.1002/dvg.20690

New DA. 1959. The adhesive properties and expansion of the chick blastoderm. J Embryol Exp Morphol 7:146–164.

New DAT. 1955. A New Technique for the Cultivation of the Chick Embryo *in vitro*. Development 3:326–331. doi:10.1242/dev.3.4.326

Nikolopoulou E, Galea GL, Rolo A, Greene NDE, Copp AJ. 2017. Neural tube closure: Cellular, molecular and biomechanical mechanisms. Development (Cambridge) 144:552–566. doi:10.1242/dev.145904

Nikolopoulou E, Hirst CS, Galea G, Venturini C, Moulding D, Marshall AR, Rolo A, de Castro SCP, Copp AJ, D.E. Greene N. 2019. Spinal neural tube closure depends on regulation of surface ectoderm identity and biomechanics by Grhl2. Nature Communications 10:1–17. doi:10.1038/s41467-019-10164-6

Oginuma M, Moncuquet P, Xiong F, Karoly E, Chal J, Guevorkian K, Pourquié O. 2017. A Gradient of Glycolytic Activity Coordinates FGF and Wnt Signaling during Elongation of the Body Axis in Amniote Embryos. Developmental Cell 40:342–353.e10. doi:10.1016/j.devcel.2017.02.001

Omana DA, Wu J. 2009. A new method of separating ovomucin from egg white. Journal of Agricultural and Food Chemistry 57:3596–3603. doi:10.1021/jf8030937

O’Rahilly R, Müller F. 2007. Neurulation in the Normal Human EmbryoCiba Foundation Symposium. John Wiley & Sons, Ltd. pp. 70–89. doi:10.1002/9780470514559.ch5

Rodler D, Sasanami T, Sinowatz F. 2012. Assembly of the inner perivitelline layer, a homolog of the mammalian zona pellucida: An immunohistochemical and ultrastructural study. Cells Tissues Organs 195:330–339. doi:10.1159/000327013

Rozbicki E, Chuai M, Karjalainen AI, Song F, Sang HM, Martin R, Knölker HJ, Macdonald MP, Weijer CJ. 2015. Myosin-II-mediated cell shape changes and cell intercalation contribute to primitive streak formation. Nature Cell Biology 17:397–408. doi:10.1038/ncb3138

Saadaoui M, Rocancourt D, Roussel J, Corson F, Gros J. 2020. A tensile ring drives tissue flows to shape the gastrulating amniote embryo. Science (1979) 367:453–458. doi:10.1126/science.aaw1965

Sadler WW, Wilgus HS, Buss EG. 1954. Incubation Factors Affecting Hatchability of Poultry Eggs. Poultry Science 33:1108–1115. doi:10.3382/ps.0331108

Sauter EA, Stadelman WJ, Harns V, McLaren BA. 1951. Methods for Measuring Yolk Index. Poultry Science 30:629–632. doi:10.3382/PS.0300629

Schmitz M, Nelemans BKA, Smit TH. 2016. A submerged filter paper sandwich for long-term Ex ovo time-lapse imaging of early chick embryos. Journal of Visualized Experiments 2016. doi:10.3791/54636

Sheng G. 2014. Day-1 chick development. Developmental Dynamics 243:357–367. doi:10.1002/DVDY.24087

Smith JL, Schoenwolf GC. 1997. Neurulation: coming to closure. Trends in Neurosciences 20:510–517. doi:10.1016/S0166-2236(97)01121-1

Spratt NT. 1947. Regression and shortening of the primitive streak in the explanted chick blastoderm. Journal of Experimental Zoology 104:69–100. doi:10.1002/jez.1401040105

Sydow H, Pieper T, Viebahn C, Tsikolia N. 2017. An early Chick embryo culture device for extended continuous observationAvian and Reptilian Developmental Biology. pp. 309–317.

Voronov DA, Taber LA. 2002. Cardiac looping in experimental conditions: Effects of extraembryonic forces. Developmental Dynamics 224:413–421. doi:10.1002/DVDY.10121

Wallingford JB, Niswander LA, Shaw GM, Finnell RH. 2013. The Continuing Challenge of Understanding, Preventing, and Treating Neural Tube Defects. Science (1979) 339:1222002–1222002. doi:10.1126/science.1222002

Xiong F, Ma W, Bénazéraf B, Mahadevan L, Pourquié O. 2020. Mechanical Coupling Coordinates the Co-elongation of Axial and Paraxial Tissues in Avian Embryos. Developmental Cell 55:354–366.e5. doi:10.1016/j.devcel.2020.08.007

Zhou J, Pal S, Maiti S, Davidson LA. 2015. Force production and mechanical accommodation during convergent extension. Development (Cambridge) 142:692–701. doi:10.1242/dev.116533

